# Analysis of Practices to Promote Reproducibility and Transparency in Anaesthesiology Research: Are Important Aspects “Hidden Behind the Drapes?”

**DOI:** 10.1101/729129

**Authors:** Ochije Okonya, Drayton Rorah, Daniel Tritz, Blake A. Umberham, Matt Wiley, Matt Vassar

## Abstract

**Introduction:** Reliable, high-quality research is essential to the field of anaesthesiology. Reproducibility and transparency has been investigated in the biomedical domain and in the social sciences, with both lacking to provide necessary information to reproduce the study findings. In this study, we investigated 14 indicators of reproducibility in anaesthesiology research.

**Methods:** We used the National Library of Medicine (NLM) catalogue to search for all anaesthesiology journals that are MEDLINE indexed and provided English texts. PubMed was searched with the list of journals to identify all publications from January 1, 2014 to December 31, 2018. We randomly sampled 300 publications that fit the inclusion criteria for our analysis. Data extraction was then conducted in a blinded, duplicate fashion using a pilot-tested Google form.

**Results:** The PubMed search of these journals identified 171,441 publications, with 28,310 being within the time frame. From the 300 publications sampled, 296 full-text publications were accessible. Most of the studies did not include materials or protocol availability statements. The majority of publications did not provide a data analysis script statement (121/122, 99% [98% to 100%]) or a preregistration statement (94/122, 77% [72% to 81%]).

**Conclusion:** Anaesthesiology research needs to drastically improve indicators of reproducibility and transparency. By making research publically available and improving accessibility to detailed study components, primary research can be reproduced in subsequent studies and help contribute to the development of new practice guidelines.

## Introduction

Reliable, high quality research is essential to the field of anaesthesiology. New research establishes the evidence base for clinical practice guidelines, modifies established protocols, updates the standard of care, affects reimbursement (with possibly significant financial implications), and informs clinical practice for anaesthesiologists. Consider the use of epidural glucocorticoid injections for spinal stenosis as an example. Now controversial, previous guidelines based upon uncontrolled trials recommended epidural steroid injections to treat spinal stenosis pain.^1–11^ These studies led to a 271% increase from 1991 to 2011 in physician use of epidural injections to treat spinal stenosis pain and a considerable cost increase from $24 million to $174 million^12^. Friedly and colleagues^12^, re-examined these recommendations and conducted a large, randomized, controlled trial that found no statistically significant difference at six weeks for epidural glucocorticoid plus local anaesthetic injection versus epidural local anaesthetic injection only. Because of the implications of research on patient care and healthcare costs, credible science should catalyse change and must be supported by reliable evidence.

The process of peer reviewing, analysing, critiquing, and, eventually, reproducing trials is the cornerstone for creating high quality, reliable, transparent, reproducible, evidence-based publications.^13^ In fact, reproducibility and transparency are core scientific principles. However, published reports may provide only limited summaries of a research study, and these published reports often fail to include key study components – raw data, detailed protocols, materials, and analysis scripts – that provide more comprehensive study details. Access to this additional information enables further analysis and verification of the conclusions from the original research.^14^ When researchers strive for transparency and allow primary research to be reproducible, we will see improved efficiency^15^, self-correction^16^, and credible published literature.^17^ Because of the vital importance of accurate research and its direct influence on patient care, publishers of high-impact journals have initiated guidelines to help improve the reproducibility and transparency of research. For example, the *British Journal of Anaesthesia* and *Anesthesia & Analgesia* provide statements in their authorship guidelines encouraging raw data to be available to readers; however, raw data is not required to be submitted for public viewing.^18,19^ Access to raw data is encouraged by these journals for statistical reproducibility^20^, additional analysis^21^, participant-level meta-analyses^22^, and the merging of future or existing datasets.^23^

Reproducibility and transparency have been assessed in biomedical and social sciences; however, practices that promote reproducibility and transparency have never been evaluated in anaesthesiology research. In this study, we queried indicators of reproducibility to assess the current climate of anaesthesiology research. Results from this investigation may be used to establish a baseline for comparison in future studies.

## Methods

This is an observational, cross-sectional study design. We used the methodology by Hardwicke and colleagues^24^ with modifications. This study did not involve human participants and was not subject to oversight by an institutional review board per the United States Code of Federal Regulations. ^25^ We report our study in accordance with guidelines for meta-epidemiological methodology research.^26^ We uploaded our protocol, data extraction form, and other necessary materials for public viewing on the Open Science Framework (https://osf.io/n4yh5/).

### Journal Selection

We used the National Library of Medicine (NLM) catalogue to search for all relevant journals using the subject terms tag Anesthesiology[ST]. This search was performed on May 29, 2019. The inclusion criteria required that journals provided full-text publications in English and were MEDLINE indexed. The list of journals in the NLM catalogue fitting the inclusion criteria were then extracted using the electronic International Standard Serial Number (ISSN) or the linking ISSN when the electronic ISSN was unavailable. This series of ISSNs were then used in a PubMed search to identify all publications within these journals. We limited the sample to publications from January 1, 2014 to December 31, 2018 then randomly sampled 300 publications that fit the inclusion criteria for our analysis (https://osf.io/7sk9m/).

### Data Extraction Training

The two investigators responsible for data extraction (OO and DR) underwent a full day of training to ensure inter-rater reliability. The training included an in-person session that reviewed the project study design, protocol, Google extraction form, and examples of where information may be contained using two sample publications. The investigators were then given three example publications from which to extract data in a blinded fashion. Following data extraction, the pair reconciled differences between them by discussion. This training session was recorded and listed online for reference (https://osf.io/tf7nw/). As a final training exercise, investigators extracted data from the first 10 publications of their sample. The investigators held a meeting to reconcile any differences in the data before extracting data from the remaining 290 publications.

### Data Extraction

Data extraction on the remaining 290 publications was then conducted in a duplicate, blinded fashion. A final consensus meeting was held with both investigators to resolve disagreements. A third investigator (DT) was available for adjudication but was not needed. We extracted data using a pilot-tested Google form based on the one provided by Hardwicke and colleagues with additions.^24^ This form queries information necessary to be reproducible, such as the availability of materials, data, protocols, or analysis scripts (https://osf.io/3nfa5/). The data extracted varied based on the study design with studies having no empirical data being excluded (e.g., editorials, commentaries [without reanalysis], simulations, news, reviews, and poems). In our Google form, we included the five-year and most-recent-year impact factor, when available. We also expanded the study design options to include: cohort studies, case series, secondary analyses, chart reviews, and cross-sectional studies. Finally, we expanded the funding options from public, private, or mixed into the more specific categories of university, hospital, public, private/industry, non-profit, or mixed.

### Evaluation of Open Access Status

We evaluated all 300 publications to see if they were freely available online through open access. We searched the Open Access Button (openaccessbutton.org) with publication titles and DOI numbers. This database actively searches for the full-text online. If the Open Access Button was unable to find the publication, authors (OO and DR) then searched Google and PubMed to determine if the full-text was available on the journal website.

### Evaluation of Replication and Whether Publications Were Included in Research Synthesis

For empirical studies, excluding meta-analysis and commentary with analysis, we searched the Web of Science to determine if the publication was cited in a replication study, meta-analysis, or systematic review. The Web of Science additionally lists information important for our study, such as the country of journal publication, five-year impact factor (when available), and most recent impact factor with the year it represents.

### Statistical Analysis

We report descriptive statistics for each of our findings with 95% confidence intervals (95% CI) using analysis functions within Microsoft Excel.

## Results

### Publication Characteristics and Availability

Our search of the NLM catalogue identified 86 anaesthesiology journals, but only 36 fit the inclusion criteria. The PubMed search of these journals identified 171,441 publications, with 28,310 being within the time-frame. From the 300 publications sampled, 296 full-text publications were obtained (296/300, 98% [95% CI: 97% to 99%]), while four only provided the abstract or could not be accessed (4/300, 1% [95% CI: 0% to 3%]). Of the 296 publications, 53% (160/300) were publicly accessible. The remaining 47% (140/300) were blocked by a paywall, which could only be accessed through academic library subscriptions (Table 1). We analyzed several anaesthesiology journals of varying five-year impact factors with a median of 2.902 [interquartile range: 1.8-4.0]. For 32 publications, the journal impact factor was unavailable. Other sample characteristics are shown in Table 1 and Supplemental Table 1.

**Table 1:** Types of Studies in Anaesthesiology.

| Table 1: Types of Studies in Anaesthesiology |  |  |
| --- | --- | --- |
| Characteristics |  | Variables |
|  |  | No. (%) |
| **Type of Study**<br>(N=296) | No empirical | 142 (48.0) |
|  | Clinical Trial | 38 (12.8) |
|  | Case study | 27 (9.1) |
|  | Laboratory | 22 (7.4) |
|  | Cohort | 21 (7.1) |
|  | Meta-analysis | 12 (4.1) |
|  | Chart Review | 10 (3.4) |
|  | Survey | 8 (2.7) |
|  | Case Series | 5 (1.7) |
|  | Commentary | 3 (1.0) |
|  | Case Control | 1 (0.3) |
|  | Multiple | 1 (0.3) |
|  | Cost effect | 0 (0.0) |
|  | Other | 6 (2.0) |
| **Study participants**<br>(N=296) | Animals | 17 (5.7) |
|  | Humans | 119 (69.9) |
|  | Both | 0 (0.0) |
|  | Neither | 160 (2.5) |
| **Country of journal publication**<br>(N=296) | US | 211 (71.3) |
|  | South Korea | 6 (2.0) |
|  | UK | 32 (10.8) |
|  | Australia | 8 (2.7) |
|  | Japan | 12 (4.1) |
|  | France | 4 (1.4) |
|  | Canada | 3 (1.0) |
|  | Italy | 8 (2.7) |
|  | India | 3 (1.0) |
|  | Poland | 7 (2.4) |
|  | Unclear | 2 (0.7) |
|  | Other | 0 (0.0) |
| **Country of first author**<br>(N=296) | US | 101 (34.1) |
|  | China | 7 (2.4) |
|  | UK | 23 (7.8) |
|  | Netherlands | 9 (3.0) |
|  | Japan | 8 (2.7) |
|  | France | 10 (3.4) |
|  | Canada | 19 (6.4) |
|  | Italy | 13 (4.4) |
|  | India | 11 (3.8) |
|  | Australia | 14 (4.7) |
|  | Unclear | 2 (0.7) |
|  | Other | 79 (26.7) |
| **5 Year Impact Factor**<br>(N=268) | Median | 2.9 |
|  | 1st quartile | 2.0 |
|  | 3rd quartile | 3.8 |
|  | Interquartile range | 1.8-4.0 |
| **Impact Factor, Most Recent Year** (N=296) | 1999 | 3 (1.0) |
|  | 2014 | 0 (0.0) |
|  | 2015 | 2 (0.7) |
|  | 2016 | 0 (0.0) |
|  | 2017 | 263 (88.9) |
|  | 2018 | 0 (0.0) |
|  | Not found | 32 (10.8) |
| **Impact Factor,**<br>**Most Recent**<br>**Year (N=268)** | Median | 3.4 |
|  | 1st quartile | 1.8 |
|  | 3rd quartile | 4.0 |
|  | Interquartile<br>range | 2.0-3.8 |

### Replication Criteria

The presence of several reproducibility criteria were analyzed, including: publication availability, conflict of interest statement, funding statement, protocol availability, raw data availability, materials availability statement, preregistration statement, and analysis script availability (Table 2). Of the 154 publications containing empirical data, 107 were assessed for a materials availability statement. Meta-analysis, case studies, case series and commentary with analysis studies were excluded from this evaluation (Figure 1). Most of the studies containing data did not include materials availability statements or protocol availability statements (104/107, 97% [95% CI: 95% to 99%]). The availability of raw data, analysis scripts, study protocol, and preregistration was accessed in 122 studies. Case studies and case series were excluded from this evaluation (Figure 1). Most of these studies did not provide a data availability statements (105/122, 86% [95% CI: 82% to 90%]). In the studies that had accessible data, only 8% included all the raw data to reproduce the study findings (1/13 [95% CI: 5% to 11%]). The majority of publications did not provide a data analysis script statement (121/122, 99% [95% CI: 98% to 100%]). Similar to analysis scripts, a majority of the publications did not contain a preregistration statement (94/122, 77% [95% CI: 72% to 81%]). Additional information is available in Table 3 and Supplemental Table 1.

**Table 2:**
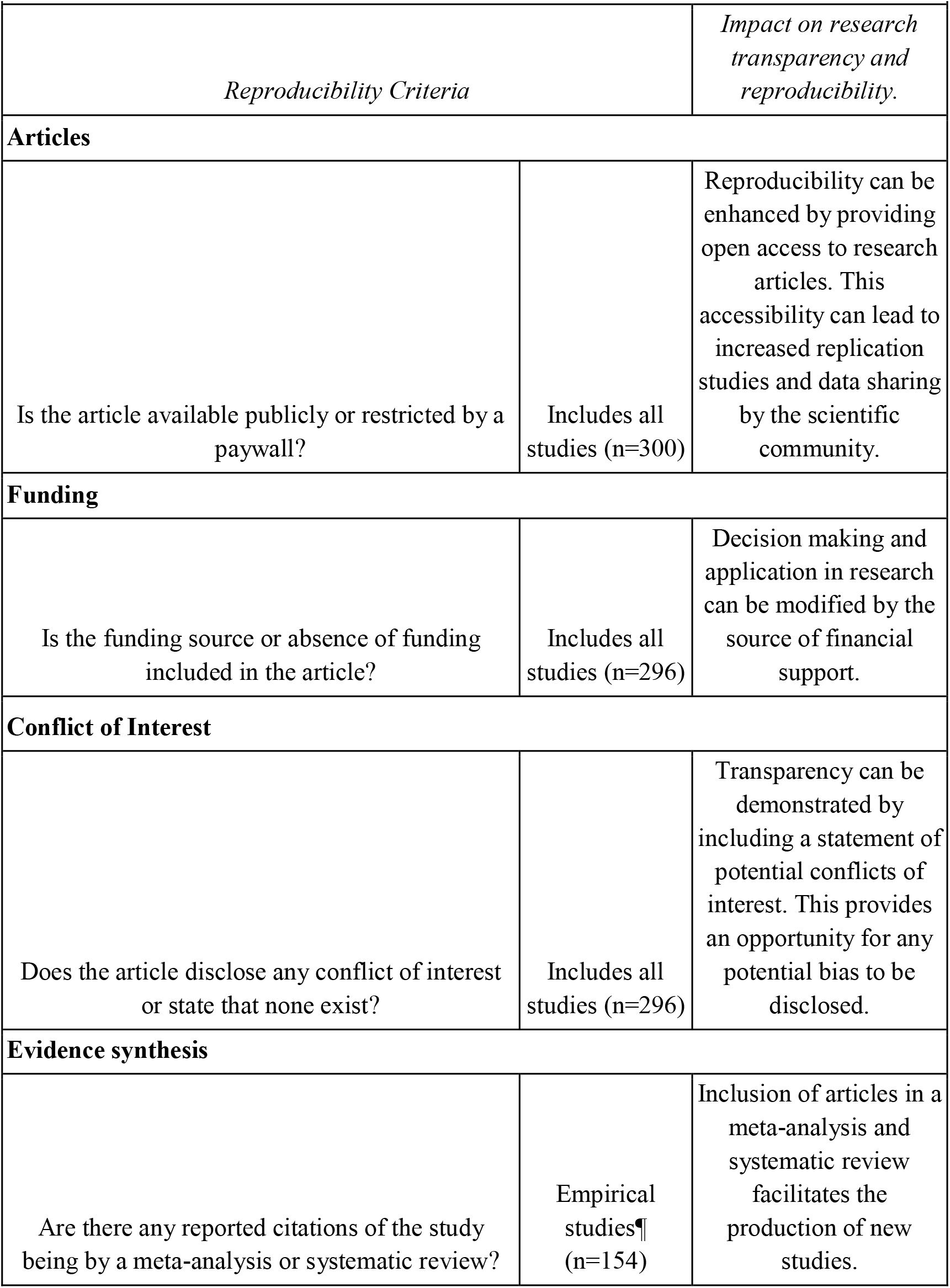
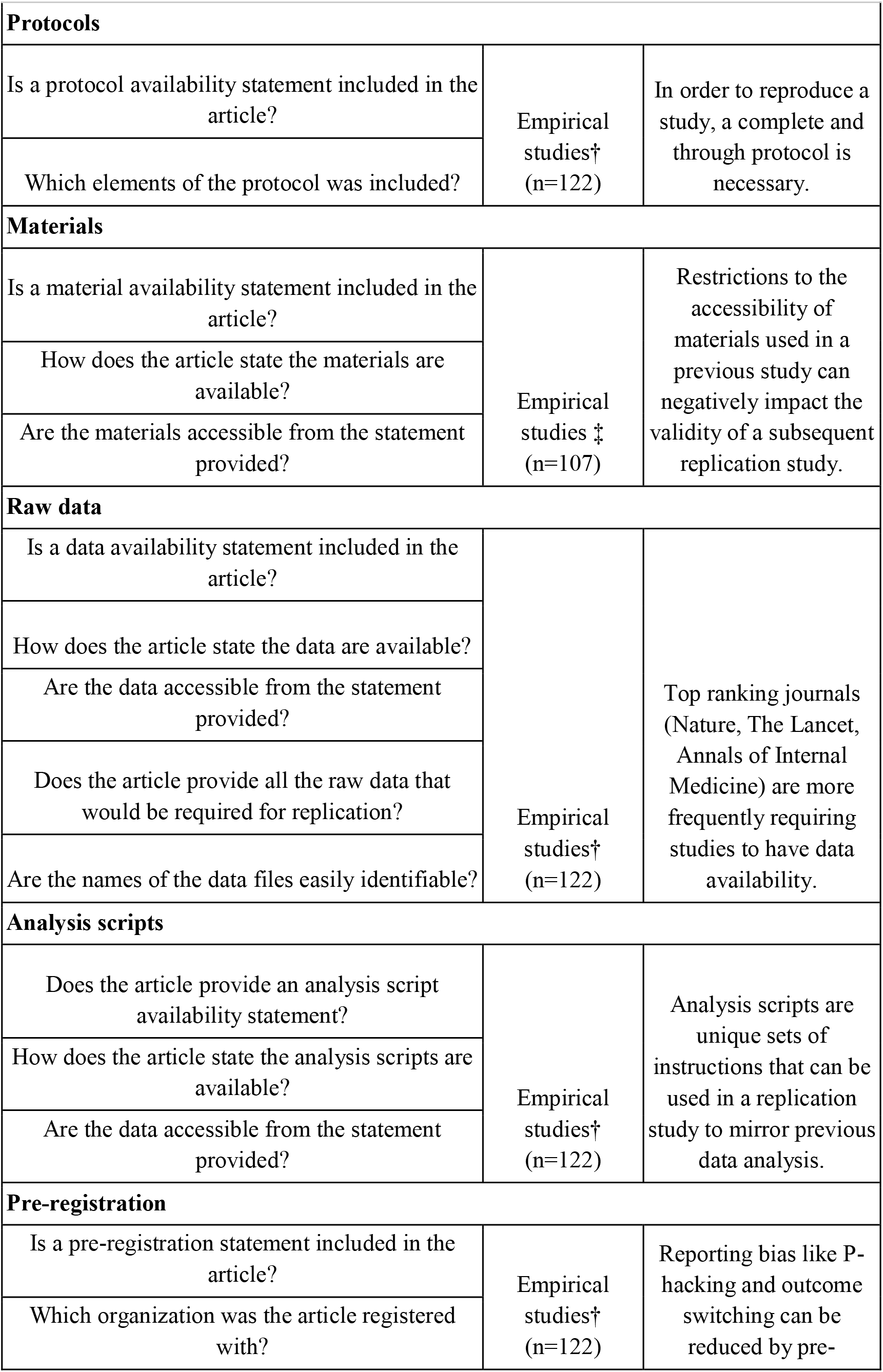
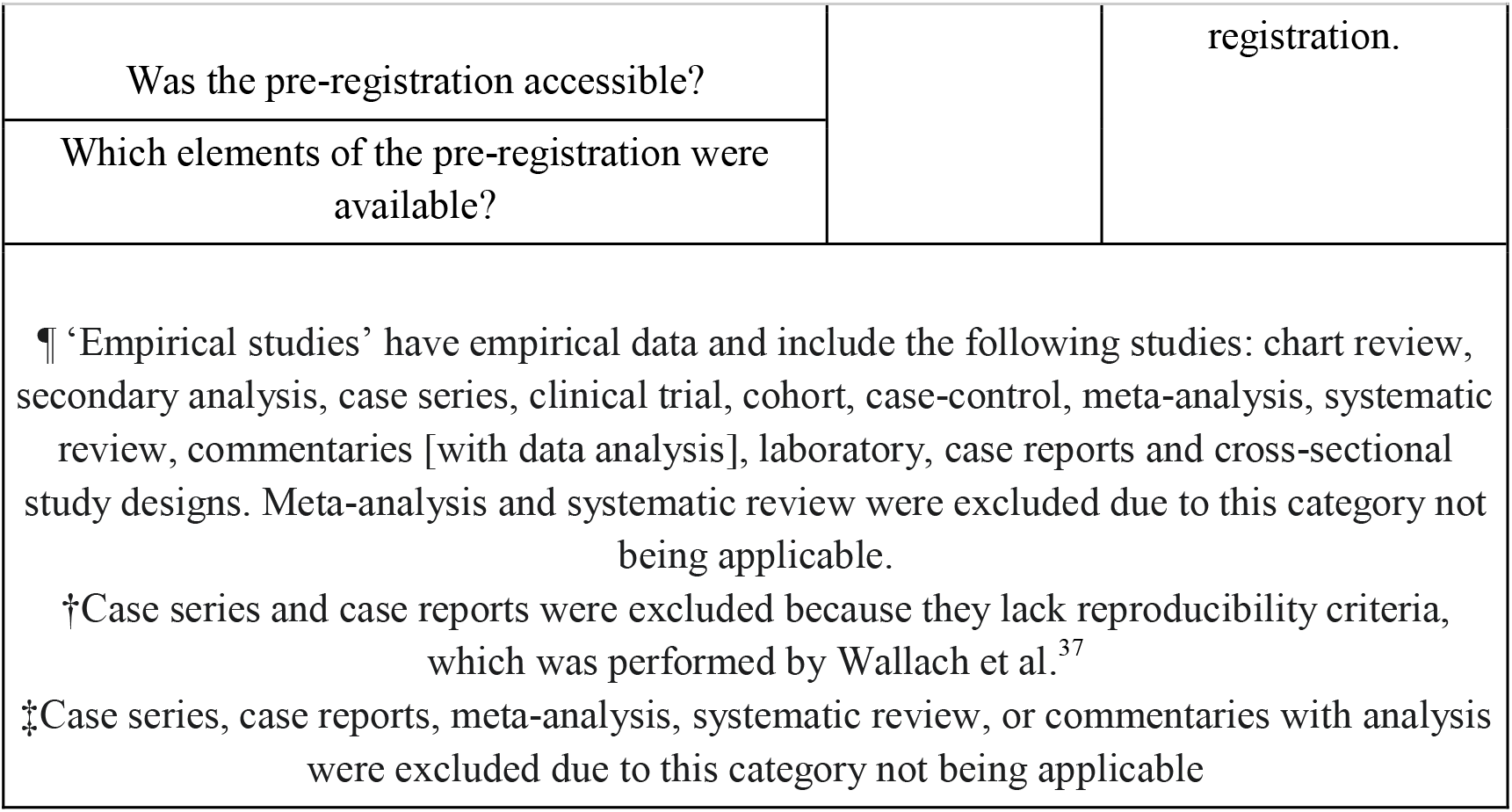
Factors Analyzed. The factors examined in each article were based upon study type. Additional information about coding and extraction is accessible at: https://osf.io/x24n3/

**Figure 1:**
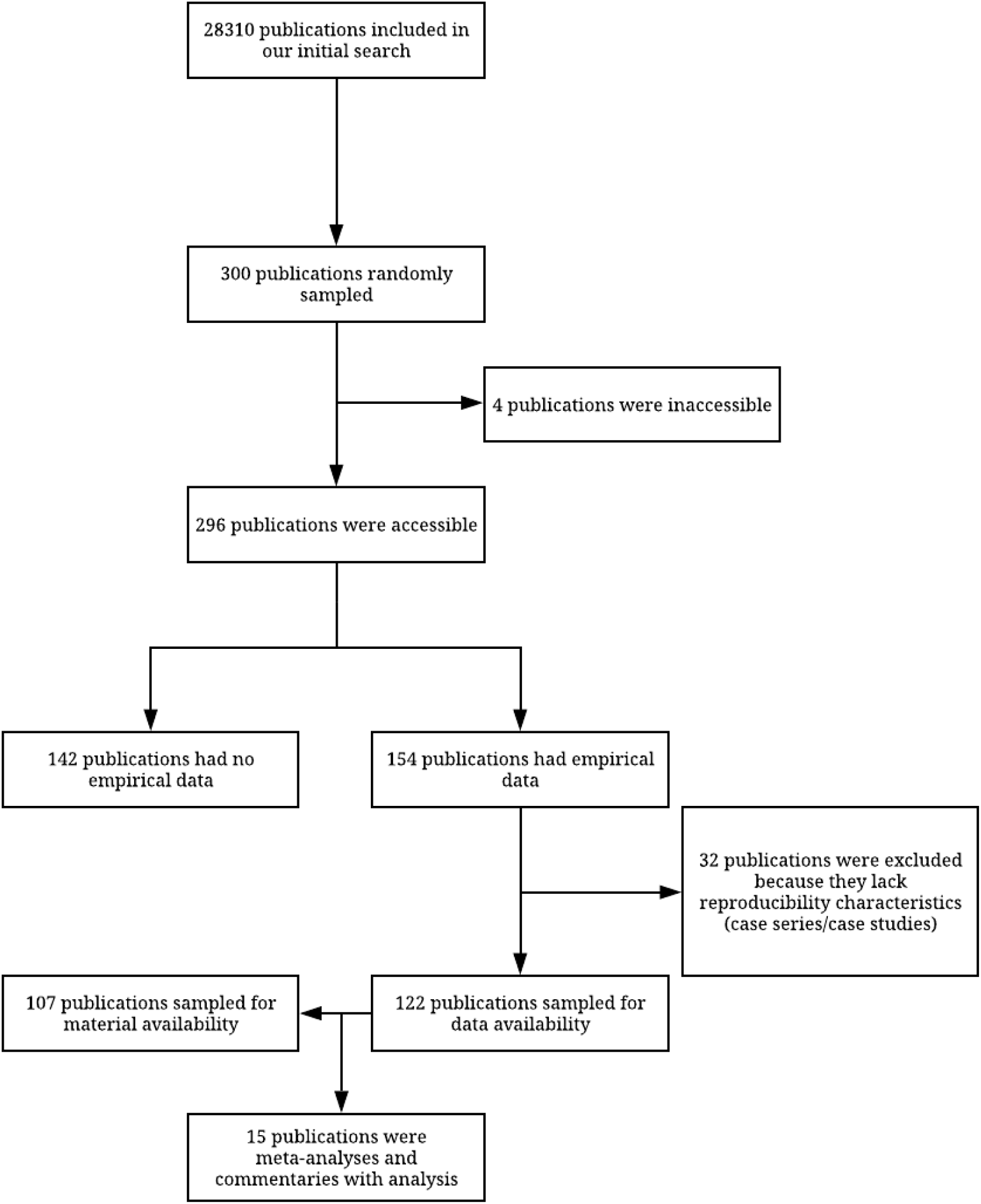
Flow diagram for inclusion and exclusion of studies

**Table 3:** Reproducibility Criteria.

| Characteristics |  | Variables |  |
| --- | --- | --- | --- |
|  |  | N (%) | 95% CI |
| **Funding (N=296)** | University | 27 (9.1) | 5.9-12.4 |
|  | Hospital | 33 (11.1) | 7.6-14.7 |
|  | Public | 15 (5.1) | 2.6-7.5 |
|  | Industry/Private | 32 (10.8) | 7.3-14.3 |
|  | Non-Profit | 8 (2.7) | 0.9-4.5 |
|  | No funding statement | 140 (47.3) | 41.6-52.9 |
|  | Not funded | 64 (21.6) | 17.0-26.3 |
|  | Mixed | 64 (21.6) | 17.0-26.3 |
| **Conflict of Interest statement (N=296)** | Conflict of interest $\geq 1$ | 35 (11.8) | 8.2-15.5 |
|  | No conflict of interest | 171 (57.8) | 52.2-63.4 |
|  | Statement not present | 90 (30.4) | 25.2-35.6 |
| **Data Availability Statement (N=122)** | Some or all data available | 16 (13.1) | 9.3-16.9 |
|  | Data not available | 1 (0.8) | 0.0-1.8 |
|  | Statement not present | 105 (86.1) | 82.1-90.0 |
| **Material Availability Statement (N=107)** | Some or all data available | 2 (18.7) | 0.3-3.4 |
|  | Materials not available | 1 (0.9) | 0.0-2.0 |
|  | Statement not present | 104 (97.2) | 95.3-99.1 |
| **Protocol Available Statement (N=122)** | Complete Protocol | 4 (3.3) | 1.3-5.3 |
|  | Statement not present | 118 (96.7) | 94.7-98.7 |
| **Analysis Scripts (N=107)** | Some or all analysis scripts available | 1 (0.8) | 0.0-1.8 |
|  | Analysis scripts not available | 0 | 0 |
|  | Statement not present | 121 (99.2) | 98.2-100 |
| **Pre-registration Statement**<br>(N=122) | Pre--registered | 28 (23.0) | 18.2-27.7 |
|  | Not pre--registration | 0 (0.0) | - |
|  | Statement not present | 94 (77.0) | 72.3-81.8 |
| **Open Access**<br>(N=300) | Publically available, Open Access Button | 81 (27.0) | 21.9-32.0 |
|  | Publically available, other means | 79 (26.3) | 21.3-31.3 |
|  | Not accessible, blocked by paywall | 140(46.7) | 41.0-52.3 |
| Abbreviations: CI, Confidence Interval. |  |  |  |

### Replication and Evidence Synthesis

The publications were analyzed for their number of citations in replication studies or systematic reviews/meta-analyses. Of the 154 publications containing empirical data, 139 studies were included in this analysis; meta-analyses, systematic reviews, and commentaries with analysis were excluded (Table 2). None of the 139 studies were cited by a replication study. Similarly, most of the publications were not cited by a systematic review or meta-analysis (122/139, 88% [95% CI: 84% to 91%]; Supplemental Table 1).

### Conflict of Interest and Funding Statements

The presence of these statements was analyzed for the accessible 296 publications. The majority of publications provided a conflict of interest statement (206/296, 70% [95% CI: 64% to 75%]), while a minority did not contain a conflict of interest statement (90/296, 30% [95% CI: 35% to 36%]). Most publications contained a funding statement (155/296, 52% [95% CI: 47% to 58%]). Of the publications containing a funding statement, 58% were funded (90/155 [95% CI: 52% to 64%]), while 42% stated no funding was received (65/155 [95% CI: 36% to 48%]). More conflict of interest statements were listed relative to funding statements. The most common sources of funding for the publications were private/industry (24/90, 27% [95% CI: 22% to 32%]) and mixed funding sources (22/90, 24% [95% CI: 19% to 29%]). Additional information is presented in Table 3.

## Discussion

To our knowledge, this is the first study attempting to objectively quantify specific indicators of reproducibility and transparency in the field of anaesthesiology. Our results show disregard for reproducibility and transparency in currently published anaesthesiology research. The majority of the publications in our sample failed to make key study components available. Materials and protocols were not routinely accessible, many authors did not provide raw data, only one publication provided an analysis script, and the majority were not preregistered. There were no published replication attempts in our sample of publications. Of the indicators we investigated, conflict of interest disclosures and funding statements, were the only indicators included by a majority of the researchers; however, there is still room for improvement. Publications failing to provide key study components can have unintended consequences when others attempt to replicate the research or when it is included in a meta-analysis or systematic review. Seitz et al. conducted a systematic review on exposure to general anaesthesia and risk of developing Alzheimer’s disease. When pooling the primary studies, only a single study specified the time duration between exposure to general anaesthesia and assessment of dementia. This lack of reporting prohibited the authors from estimating a pooled effect estimate for this important outcome. ^27^ Had better reporting been performed by the primary study authors, this analysis would have been possible.

The lack of publicly available protocols, materials, and data in anaesthesiology literature is concerning. These research methodologies allow for independent verification of results and for ensuring that researchers actually did what they planned to do. For example, comparing study protocols or preregistrations with published reports allows for the evaluation of outcome switching or selective reporting bias. This bias occurs when study authors add, remove, promote, or demote outcomes in a study based on whether these outcomes were statistically significant. This form of bias can be problematic, leading to misinterpretations of clinical trials. Several studies have examined the pervasiveness of this problem in the medical literature. ^28,29^ P-hacking, the practice of running multiple statistical analyses until statistical significance is achieved, is another significant problem that can be mitigated if statistical analysis plans are transparent and available for evaluation. HARKing (Hypothesizing After Results are Known) occurs when post-hoc findings are insinuated to be *a priori* hypothesis-driven analyses in published study reports. Thus, all three forms of research malpractice may be inspected if study authors provide all available materials and preregister their studies.

### Future for Reproducibility and Transparency

Improving the reproducibility crisis in science requires an actionable response by multiple stakeholder groups. Below, we outline recommendations being adopted inside and outside of medicine that may be useful to the field of anaesthesiology. Here, we focus on the role of academic journals and funders, although, certainly, the researchers themselves, peer reviewers, institutional review boards, and others play a role in this improvement.

In a recent article, Adams describes efforts by the *British Journal of Anaesthesia* to improve study reproducibility by considering reproducibility beyond the methods and results. ^30^ The journal’s editor-in-chief has created a novel approach for arriving at more accurate conclusions by involving independent reviewers to write discussions and conclusions during the peer review process when provided with the submitter’s raw data. The idea would attempt to eliminate the original authors’ conflicts of interests and allegiance biases. These biases can alter the interpretation of their results. Although we did not inquire about reproducibility with regards to drawing conclusions from submitted data and methodology, this seems to be an additional measure journals could consider taking in order to ensure published material is not misconstrued. The journal *Anesthesiology* uses custom software designed to evaluate a study’s adherence to reporting guidelines like the Consolidated Standards for Reporting Trials (CONSORT) and Animal Research: Reporting of *In Vivo* Experiments (ARRIVE). Outside of medical research, novel approaches are also being explored. *The American Journal of Political Science* requires manuscripts accepted for publication to provide sufficient materials in the text and supporting materials for independent researchers to verify the analytic results. Upon submission of the final draft, these materials are verified that they do, in fact, reproduce all results in the manuscript by an independent statistical group at a university. Following this confirmation process, the dataset is deposited online. Thus, publication in the journal is contingent upon authors providing all necessary materials and successful verification of all data.

The future of reproducible research does not rest solely on the shoulders of academic journals. Funders play an important role, too. The NIH and NSF have both developed processes to improve the reproducibility of studies funded by federal tax dollars. ^31,32^ The Wellcome Trust is a “politically and financially independent” group that funds thousands of researchers internationally. The Trust influences researchers and policy makers to improve the methodological quality of publications. In order to receive funding from this group, authors are expected to include accurate records of the methods, procedures, and approvals so that the findings can be replicated. ^33^ The Bill and Melinda Gates Foundation is also a significant funding source for researchers. Although they do not include as rigorous of guidelines on manuscript submission, they do require that all publications funded by them be immediately open access. The push for open access aids in the dissemination of new research and findings across the world and can actively change the direction of subsequent research designs. ^34^ Both of these foundations emphasise important aspects of transparent and reproducible research.

Our study has both strengths and limitations. Concerning its strengths, our study examined a wide range of anaesthesiology publications published across several journals. The random sample of these publications used in this study should improve the generalizability of our findings. We used double data extraction throughout the data collection process. This form of data extraction, which incorporates two coders who are blinded to the decision making of the other, is considered the gold standard by the systematic review community and is advocated by the Cochrane Collaboration. ^35^ Additionally, we have provided our study protocol, data, and other pertinent materials to improve the reproducibility and transparency of this study. Regarding its limitations, our data collection was sampled from publications dated from 2014 to 2018 and is meant to be a general overview rather than a complete analysis of anaesthesiology publications. Our data collection is also limited to publications in the field of anaesthesiology. We recommend investigating reproducibility and transparency in other fields of medicine as there is often overlap which can contribute to the development of clinical guidelines and protocols. For example, the recent Enhanced Recovery After Surgery (ERAS) protocol developed for Cardiac Surgery published in *JAMA Surgery* included several randomized control trials and meta-analyses that would not necessarily have been found in specific anaesthesiology journals.^36^

In conclusion, anaesthesiology research needs to drastically improve with regards to reproducibility and transparency. This analysis is consistent with previous studies in the biomedical and social science research. We speculate our findings are also consistent in other fields of medicine; however, we recommend further analysis in order to catalyse change in those fields. Our goal of this study is to offer a foundation for publishers to consider when evaluating the validity of a study and for authors and researchers to consider when developing their primary research projects. By including these indicators in primary research, anaesthesiology publications can become more valid, transparent, and reproducible. By making research easily accessible online and by improving accessibility to the detailed study components (raw data, materials, protocols, and analysis scripts) primary research can be reproduced in subsequent studies and help contribute to the development of new practice guidelines, helping change patient care through evidence-based conclusions.

## Authors’ Contributions

1. Substantial contributions to the conception or design of the work; or the acquisition, analysis, or interpretation of data for the work: OO, DR, DT, BU, MW, MV
2. Drafting the work or revising it critically for important intellectual content: OO, DR, DT, BU, MW, MV
3. Final approval of the version to be published: OO, DR, DT, BU, MW, MV

## Acknowledgements

None.

---

**Supplemental Table 1:** Additional Reproducibility Criteria.

| Supplemental Table 1: Additional Reproducibility Criteria |  |  |
| --- | --- | --- |
| Characteristics |  | N |
| Material availability | Institutional or Personal | 0 |
|  | Material hosted by the journal | 2 |
|  | Third party website | 0 |
|  | Only available upon request | 0 |
|  | Yes, accessible | 1 |
|  | No, not accessible | 1 |
| Data availability | Institutional or Personal | 3 |
|  | Data hosted by the journal | 11 |
|  | Third party website | 1 |
|  | Only available upon request | 1 |
|  | Other | 0 |
|  | Yes, available for download | 13 |
|  | No, not available for download | 3 |
|  | Yes, files are clearly labeled | 7 |
|  | No, files are not clearly labeled | 8 |
|  | Yes, all raw data contained in files | 1 |
|  | No, all raw data not contained in files | 12 |
|  | Raw data availability unclear | 2 |
| Analysis Script | Institutional or Personal | 0 |
|  | Analysis Script hosted by the journal | 0 |
|  | Third party website | 1 |
|  | Only available upon request | 0 |
|  | Other | 0 |
|  | Yes, accessible and available for download | 0 |
|  | No, not accessible and available for download | 1 |
| **Pre-registration** | Pre-registered, ClinicalTrials.gov | 15 |
|  | Other | 13 |
|  | Yes, accessible | 22 |
|  | No, not accessible | 0 |
|  | Pre-registered hypothesis | 4 |
|  | Pre-registered methods | 17 |
|  | Pre-registered analysis | 13 |
| **Protocol** | Hypotheses was included in the protocol | 0 |
|  | Methods were included in the protocol | 4 |
|  | Analysis plan was included in the protocol | 0 |
| **Replication Studies (N=139)** | Novel study | 139 (100.0) |
|  | Replication | 0 (0.0) |
| **Cited by Systematic Review/Meta-Analysis (N=139)** | No Citations | 122 (87.8) |
|  | A Single Citation | 11 (7.9) |
|  | One to Five Citations | 6 (4.3) |
|  | Greater than Five Citations | 0 (0.0) |

## References

1. Van Boxem K, Rijsdijk M, Hans G, de Jong J, Kallewaard JW, Vissers K, et al. Safe Use of Epidural Corticosteroid Injections: Recommendations of the WIP Benelux Work Group. Pain Pract. 2019 Jan;19(1):61–92.

2. Bagley C, MacAllister M, Dosselman L, Moreno J, Aoun SG, El Ahmadieh TY. Current concepts and recent advances in understanding and managing lumbar spine stenosis. F1000Res [Internet]. 2019 Jan 31;8. Available from: http://dx.doi.org/10.12688/f1000research.16082.1

3. Cohen SP, Wallace M, Rauck RL, Stacey BR. Unique aspects of clinical trials of invasive therapies for chronic pain. Pain Reports. 2019;4(3):e687.

4. Kim J. A Retrospective Study on Combined Traditional Korean Medicine Treatment of Cervical Radiculopathy Patients Who Underwent Ineffective Epidural Steroid Injection Treatment [Internet]. Vol. 35, Journal of Acupuncture Research. 2018. p. 248–51. Available from: http://dx.doi.org/10.13045/jar.2018.00248

5. Lewandrowski K-U. Successful outcome after outpatient transforaminal decompression for lumbar foraminal and lateral recess stenosis: The positive predictive value of diagnostic epidural steroid injection [Internet]. Vol. 173, Clinical Neurology and Neurosurgery. 2018. p. 38–45. Available from: http://dx.doi.org/10.1016/j.clineuro.2018.07.015

6. Shehab N, Brown MN, Kallen AJ, Perz JF. U.S. Compounding Pharmacy-Related Outbreaks, 2001–2013 [Internet]. Vol. 14, Journal of Patient Safety. 2018. p. 164–73. Available from: http://dx.doi.org/10.1097/pts.0000000000000188

7. Chuan Yen T, Mohler J, Dohm M, Laksari K, Najafi B, Toosizadeh N. The Effect of Pain Relief on Daily Physical Activity: In-Home Objective Physical Activity Assessment in Chronic Low Back Pain Patients after Paravertebral Spinal Block. Sensors [Internet]. 2018 Sep 12;18(9). Available from: http://dx.doi.org/10.3390/s18093048

8. Lafian AM, Torralba KD. Lumbar Spinal Stenosis in Older Adults. Rheum Dis Clin North Am. 2018 Aug;44(3):501–12.

9. Jamison DE, Cohen SP. Critically Evaluating the Evidence for Epidural Injections for Failed Back Surgery Syndrome: Should Pain Physicians Be Bracing for Impact? Pain Med [Internet]. 2018 May 24; Available from: http://dx.doi.org/10.1093/pm/pny101

10. Nelson AM, Nagpal G. Interventional Approaches to Low Back Pain [Internet]. Vol. 31, Clinical Spine Surgery. 2018. p. 188–96. Available from: http://dx.doi.org/10.1097/bsd.0000000000000542

11. Olafsen NP, Herring SA, Orchard JW. Injectable Corticosteroids in Sport [Internet]. Clinical Journal of Sport Medicine. 2018. p. 1. Available from: http://dx.doi.org/10.1097/jsm.0000000000000603

12. Friedly JL, Comstock BA, Turner JA, Heagerty PJ, Deyo RA, Sullivan SD, et al. A randomized trial of epidural glucocorticoid injections for spinal stenosis. N Engl J Med. 2014 Jul 3;371(1):11–21.

13. Munafò MR, Nosek BA, Bishop DVM, Button KS, Chambers CD, Percie du Sert N, et al. A manifesto for reproducible science. Nat hum behav. 2017 Jan 10;1(1):0021.

14. Klein O, Hardwicke TE, Aust F, Breuer J, Danielsson H, Mohr AH, et al. A practical guide for transparency in psychological science. Collabra: Psychology [Internet]. 2018;4(1). Available from: https://www.collabra.org/articles/10.1525/collabra.158/?utm_source=TrendMD&utm_medium=cpc&utm_campaign=Collabra%253A_Psychology_TrendMD_0

15. Chalmers I, Glasziou P. Avoidable waste in the production and reporting of research evidence. Obstet Gynecol. 2009 Dec;114(6):1341–5.

16. Ioannidis JPA. Why Science Is Not Necessarily Self-Correcting. Perspect Psychol Sci. 2012 Nov;7(6):645–54.

17. Vazire S. Quality uncertainty erodes trust in science. Collabra: Psychology [Internet]. 2017;3(1). Available from: https://collabra.org/articles/10.1525/collabra.74/print/

18. Editorial Manager - Anesthesia & Analgesia [Internet]. [cited 2019 Jun 20]. Available from: http://edmgr.ovid.com/aa/accounts/ifauth.htm

19. Elsevier. Guide for authors - British Journal of Anaesthesia - ISSN 0007-0912 [Internet]. [cited 2019 Jun 20]. Available from: https://www.elsevier.com/journals/british-journal-of-anaesthesia/0007-0912/guide-for-authors

20. Hardwicke TE, Mathur MB, MacDonald KE, Nilsonne G, Banks GC, Kidwell M, et al. Data availability, reusability, and analytic reproducibility: Evaluating the impact of a mandatory open data policy at the journal Cognition [Internet]. Available from: http://dx.doi.org/10.31222/osf.io/39cfb

21. Steegen S, Tuerlinckx F, Gelman A, Vanpaemel W. Increasing Transparency Through a Multiverse Analysis [Internet]. Vol. 11, Perspectives on Psychological Science. 2016. p. 702–12. Available from: http://dx.doi.org/10.1177/1745691616658637

22. Tierney JF, Vale C, Riley R, Smith CT, Stewart L, Clarke M, et al. Individual Participant Data (IPD) Meta-analyses of Randomised Controlled Trials: Guidance on Their Use. PLoS Med. 2015 Jul;12(7):e1001855.

23. Voytek B. The Virtuous Cycle of a Data Ecosystem [Internet]. Vol. 12, PLOS Computational Biology. 2016. p. e1005037. Available from: http://dx.doi.org/10.1371/journal.pcbi.1005037

24. Hardwicke TE, Wallach JD, Kidwell M, Ioannidis J. An empirical assessment of transparency and reproducibility-related research practices in the social sciences (2014-2017) [Internet]. 2019. Available from: http://dx.doi.org/10.31222/osf.io/6uhg5

25. eCFR — Code of Federal Regulations [Internet]. [cited 2019 Jun 27]. Available from: https://www.ecfr.gov/cgi-bin/retrieveECFR?gp=&SID=83cd09e1c0f5c6937cd9d7513160fc3f&pitd=20180719&n=pt45.1.46&r=PART&ty=HTML

26. Murad MH, Wang Z. Guidelines for reporting meta-epidemiological methodology research. Evid Based Med. 2017 Aug;22(4):139–42.

27. Seitz DP, Shah PS, Herrmann N, Beyene J, Siddiqui N. Exposure to general anesthesia and risk of Alzheimer’s disease: a systematic review and meta-analysis. BMC Geriatr. 2011 Dec 14;11:83.

28. Mathieu S, Boutron I, Moher D, Altman DG, Ravaud P. Comparison of registered and published primary outcomes in randomized controlled trials. JAMA. 2009 Sep 2;302(9):977–84.

29. Rankin J, Ross A, Baker J, O’Brien M, Scheckel C, Vassar M. Selective outcome reporting in obesity clinical trials: a cross-sectional review. Clin Obes. 2017;7(4):245–54.

30. Adam D. Reproducibility trial publishes two conclusions for one paper. Nature. 2019 Jun;570(7759):16.

31. The Institute of Education Sciences and The National Science Foundation. Common Guidelines for Education Research and Development [Internet]. 2013 [cited 2019 Jun 28]. Available from: https://www.nsf.gov/pubs/2013/nsf13126/nsf13126.pdf

32. Rigor and Reproducibility [Internet]. National Institutes of Health (NIH). [cited 2019 Jun 28]. Available from: https://www.nih.gov/research-training/rigor-reproducibility

33. Good research practice guidelines | Wellcome [Internet]. [cited 2019 Jun 28]. Available from: https://wellcome.ac.uk/funding/guidance/good-research-practice-guidelines

34. Glynn JR, Thomas SL. Open access policy. Lancet. 2013 Jun 15;381(9883):2082; discussion 2082.

35. Higgins JPT, Green S. Cochrane Handbook for Systematic Reviews of Interventions. John Wiley & Sons; 2011. 672 p.

36. Engelman DT, Ben Ali W, Williams JB, Perrault LP, Reddy VS, Arora RC, et al. Guidelines for Perioperative Care in Cardiac Surgery: Enhanced Recovery After Surgery Society Recommendations. JAMA Surg [Internet]. 2019 May 4; Available from: http://dx.doi.org/10.1001/jamasurg.2019.1153

37. Wallach JD, Boyack KW, Ioannidis JPA. Reproducible research practices, transparency, and open access data in the biomedical literature, 2015-2017. PLoS Biol. 2018 Nov;16(11):e2006930.

